# A Bioinformatics Pipeline for Whole Exome Sequencing: Overview of the Processing and Steps from Raw Data to Downstream Analysis

**DOI:** 10.1101/201145

**Authors:** Narendra Meena, Praveen Mathur, Krishna Mohan Medicherla, Prashanth Suravajhala

## Abstract

Recent advances in next generation sequencing (NGS) technologies have given an impetus to find causality for rare genetic disorders. Since 2005 and aftermath of the human genome project, efforts have been made to understand the rare variants of genetic disorders. Benchmarking the bioinformatics pipeline for whole exome sequencing (WES) has always been a challenge. In this protocol, we discuss detailed steps from quality check to analysis of the variants using a WES pipeline comparing them with reposited public NGS data and survey different techniques, algorithms and software tools used during each step. We observed that variant calling performed on exome and whole genome datasets have different metrics generated when compared to variant callers, GATK and VarScan with different parameters. Furthermore, we found that VarScan with strict parameters could recover 80-85% of high quality GATK SNPs with decreased sensitivity from NGS data. We believe our protocol in the form of pipeline can be used by researchers interested in performing WES analysis for genetic diseases and by large any clinical phenotypes.

## INTRODUCTION

Next Generation Sequencing (NGS) technologies have paved the way for rapid sequencing efforts to analyze a wide number of samples. From the whole genome to transcriptome to exome, it has changed the way we look at nonspecific germline variants, somatic mutations, structural variant besides identifying associations between a variant and human genetic disease (Singleton et al. 2011). This can help understand the complex genetic disorders to get better diagnosis and assess disease risk. The analysis of exome sequencing data to find variants, however still poses multiple challenges. For example, there are several commercial and open source pipelines but configuring them in terms of benchmarking and optimizing them is a time consuming process (Pabinger et al. 2012; Guo et al 2015). Among the steps, *viz*. quality check, alignment, recalibration, variant calling, variant annotation, one needs to reach consensus on the set of tools following which one's output should be fed as other tool’s input (Gentleman et al. 2004, Stajich et al. 2002, Chang et al. 2012). While integrating, it would be appropriate to check and use the tools before finally reproducing and maintaining highly heterogeneous pipelines (Hwang et al. 2015). In this protocol, we discuss the steps for whole exome sequence (WES) analyses and it’s pipeline to identify variants from exome sequence data. Our pipeline includes open source tools that include a number of tools from quality check to variant calling (see *Supplementary information*).

## MATERIALS

The raw file (fastq) is subjected to different steps such as quality check, indexing, alignment, sorting, duplication removal, variant calling, variant annotation and finally downstream bioinformatics annotation (Pabinger et al. 2014)(Figure 1). It integrates bowtie2 (Langmead et al. 2012), samtools (Li et al. 2009), FastQC (Andrews, 2010), VarScan (Koboldt et al. 2012) and bcftools (Li et al. 2009), apart from necessary files containing the human genome (Venter et al. 2001), alignment indices (Trapnell, 2009), known variant databases (Landrum et al. 2014, Sherry et al. 2001, Auton et al. 2015). Keeping in view of the fact that the benchmark metrics for pipelines is an essential step, we have ensured that our pipeline is benchmarked on a sample fastq file taken from a human genome project. As the pipeline runs on Linux, all commands are case sensitive wherever used. Whereas this pipeline was run on a 64GB RAM with 8 core CPUs in an Ubuntu operating system (14.04 LTS machine), this can be run on a minimum 16 GB RAM machine based on the size of raw fastq file. However, for benchmarking the datasets was done on a 1 TB RAM with 32 cores (Dell) machine. A shell script (with an extension sh) was created with all the commands as detailed below.

**Figure 1:**
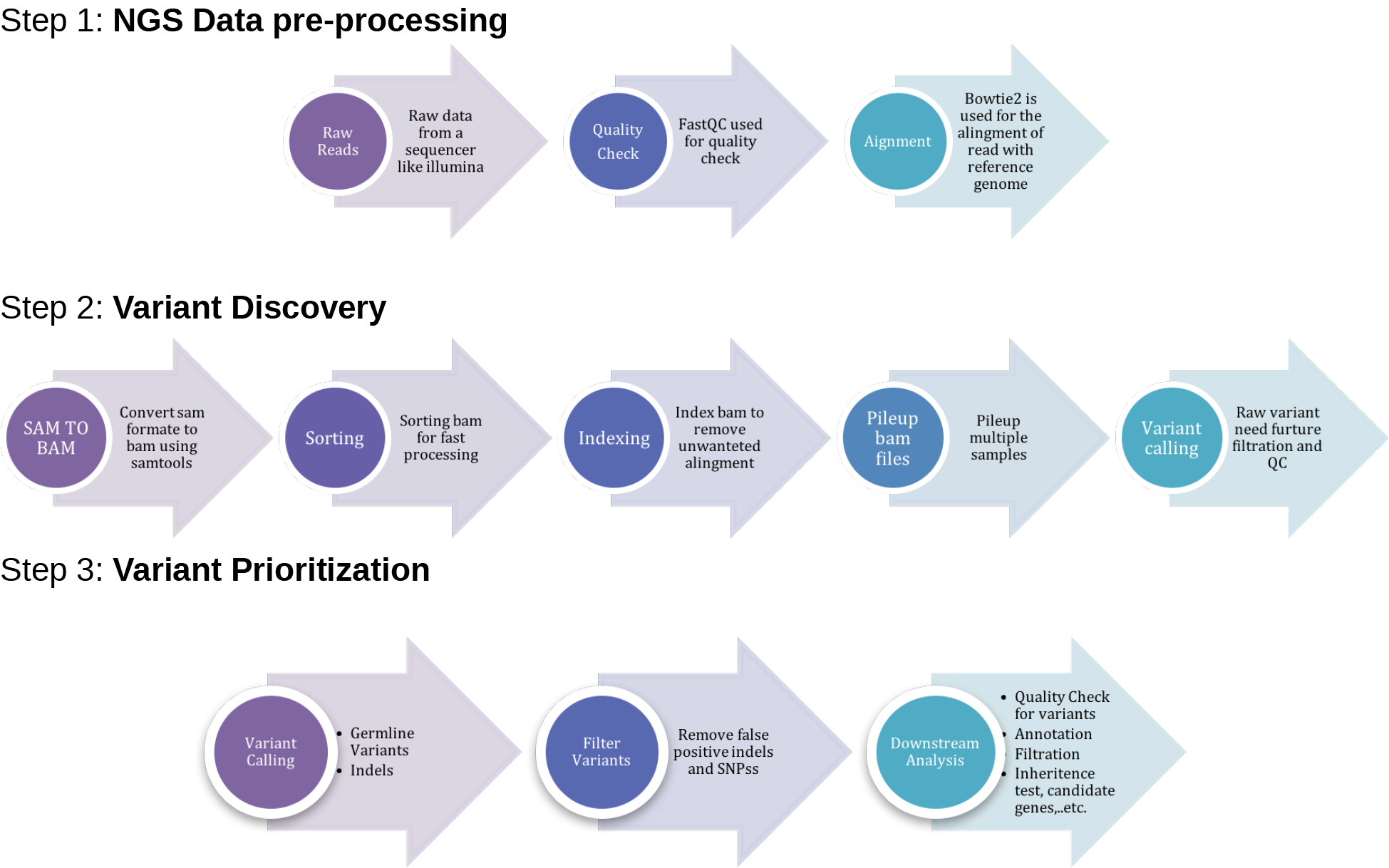
The pipeline involving three important phases, *viz*. preprocessing, variant discovery and prioritization of variants.

## PROCEDURE

### (1) Preprocessing the raw data Quality check

NGS data analysis depends on the raw data control as it provides a quick insight into the quality of the sequences. This will potentially reduce the amount of further downstream analyses with early identification of questionable samples. The ideal base quality scores for Phred (Cock et al. 2010) have paved way for the best quality scores for GC content (ca. 50% threshold) and the nucleotide distribution across all reads. In our pipeline, we used FastQC (with default Phred = 20 value) as it plots the read depth and quality score besides a host of other statistical inferences.

(i) ./fastqc ~/samples/sample1.fastq

FastQC generates an HTML formatted report with box plots and graph plots for mean quality scores for sequences, read length and depth along with the intended coverage (See Figure 2)

**Figure 2:**
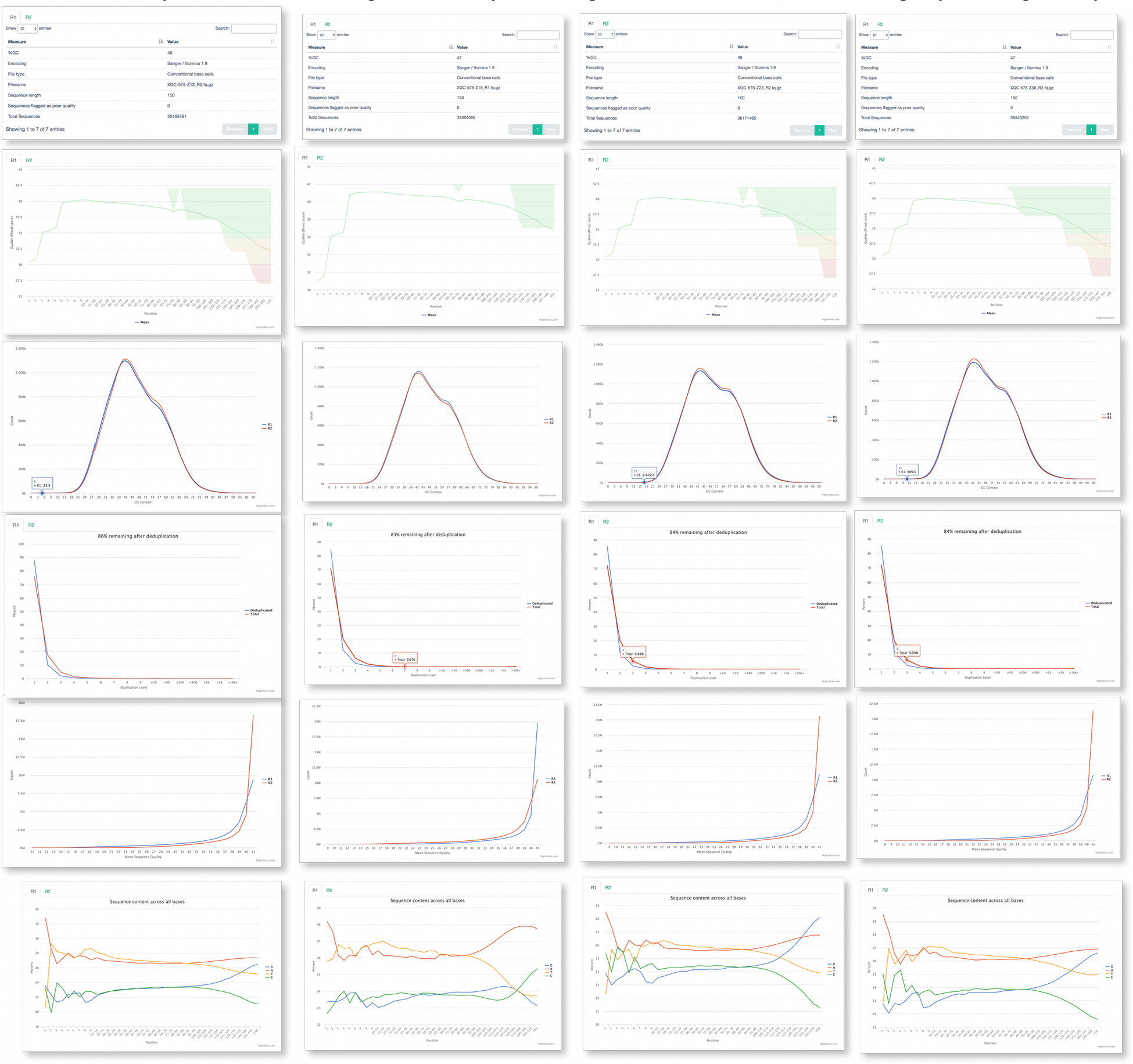
A pictorial representation containing the box plots and figures of FASTQC run containing information on statistics, quality, read coverage, depth, yield, based per read call etc.

**Table 1:**
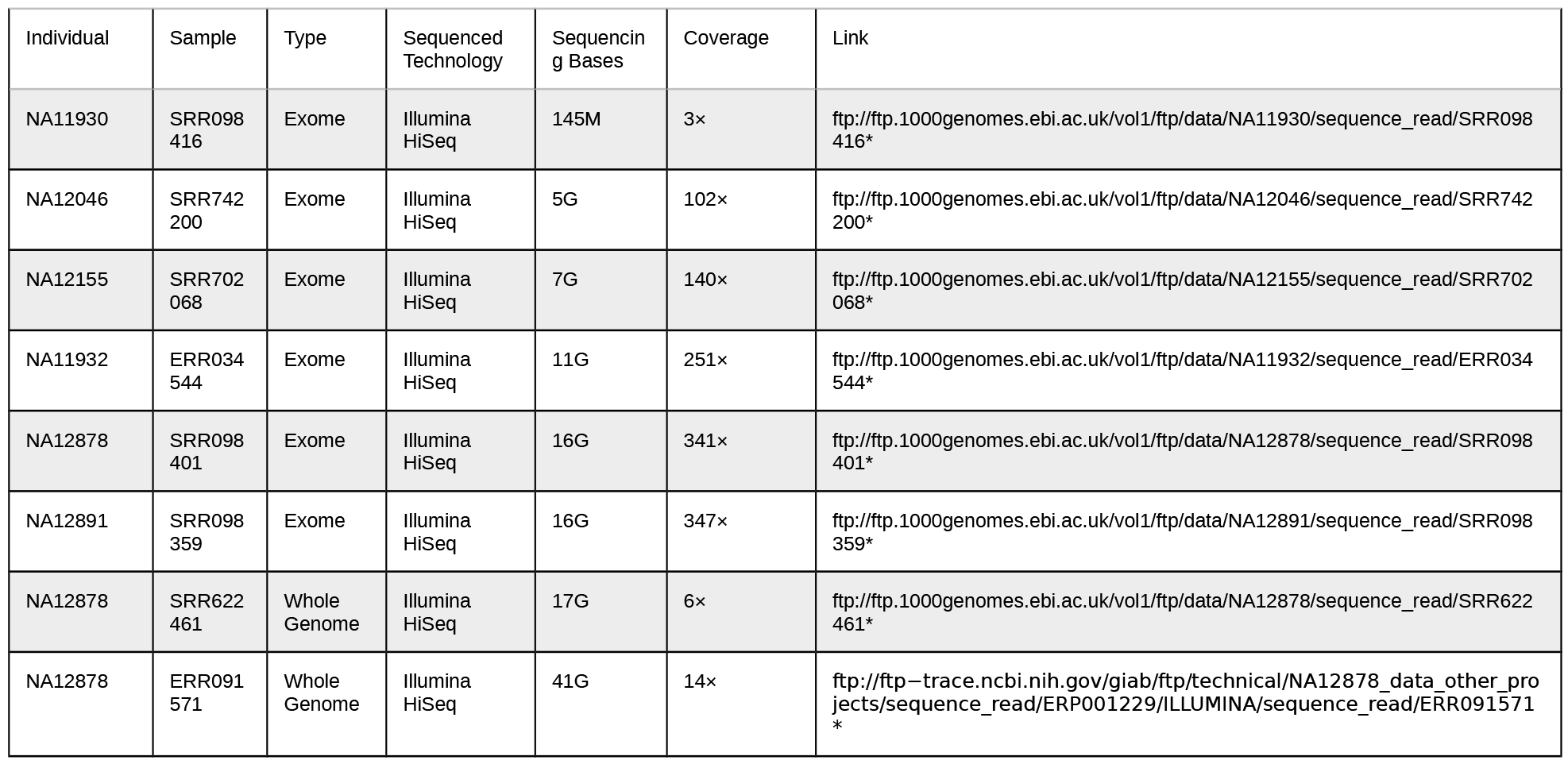
1000 Genomes samples used in benchmarking

### Indexing human genome using bowtie2

Bowtie2-build is used to index reference genome which works at high speed and memory efficient way.

(ii) *./bowtie2-build –u 10 indexes/references/reference.fq reference*

When the command is expedited, the current directory would contain four new files ending with suffices .1.bt2, .2.bt2, .3.bt2, .4.bt2, .rev.1.bt2, and .rev.2.bt2. While the first four files are the forward strands, the rev files indicate the reverse indexed sequences.

### Alignment and post processing

Bowtie2 is used for short read alignment. What makes bowtie2 interesting is the use of very little RAM with accuracy and modest performance in indexing the alignment (Langmead et al. 2012). The mismatch or any sequencing errors or small genetic variation between samples and reference genome could be checked using the following command:

(iii) *./bowtie2 -x reference_filename -1 path/filename1 -2 path/filename2 > filename.sam (The -2 option may be omitted for single-end sequences)*

Bowtie2 aligns a set of unpaired reads(in fastq or .fq format) to the reference genome using the Ferragina and Manzini (FM)-index (Langmead et al. 2012). The alignment results output in SAM format (Li et al. 2009) and a short alignment summary is written to the console.

**Samtools** is a collection of tools to manipulate the resulting alignment in SAM/BAM format. SAM stands for sequence alignment/map format and it’s corresponding format is binary mapped format (BAM). SAM is converted into different alignment formats, sorted, merged, aligned or duplicates removed and finally SNPs and short indel variants are called; whereas the BAM and following indices (.bai) are used to view the index of the binary aligned sequences. The basic need for having the binary mapped files is to save the memory.

(iv) *./samtools view -bS sample1.sam > sample1.bam*

### Sorting BAM

A sorted BAM file is used for streamlined data processing to avoid extra alignment when loaded into memory. Whereas it can be easily indexed for faster retrieval of alignment, overlapping is needed to retrieve alignment from a region.

(v) *~/samtools sort sample1.bam sample1.sorted*

*Samtools sort* is used to convert the BAM file to a sorted BAM file and *samtools index* to index BAM file.

(vi) *~/samtools index sample1.sorted.bam*

### Pileup all samples

*Samtools mpileup* step is used to analyze multiple samples across all samples thus giving coverage to all mappable reads.

(vii) ~/samtools mpileup -E -uf reference.fa sample1.bam > sample1.mpileup

### (2) Variant calling

To call variants from NGS data, VarScan among other tools provide heuristic statistical approaches, that give the desired threshold for reading depth, base quality, variant allele frequency and statistical confidence over other bayesian methods. VarScan uses *SAMtools mpileup* data as input and there are a number of options included for variant calling. For each position, the variants, which doesn’t meet the user input criteria of coverage, number of reads, variant alleles frequency and Fisher’s exact test, P-value are filtered out. This step is a prerequisite to identify those candidate mutations underlying the phenotype/disease.

### Germline variants

For germline variants, mutations that an individual inherits from their parents, or SNV calling VarScan mpileup2snp protocol is used.

(viii) *java –jar VarScan.jar mpileup2snp sample.mpileup >sample.VarScan.snp*

### Indel calling

Detecting of insertion and deletion (Indels) is the second most abundant source of finding genetic mutations in human population in a reliable manner. VarScan mpileup2indels protocol is used to call indels. The sensitivity and range for calling indels are determined by the respective alignments.

(xi) *java –jar VarScan.jar mpileup2indel sample.mpileup >sample.VarScan.indel*

### Variant filter

To get rid of false variants call and remove overlapping between SNPs and indels, a filtering option is applied on the resultant variant calling files which provide SNV and indels with higher confidence. An option to generate readcount report can also be used with VarScan.

(x) *java –jar VarScan.jar filter sample.VarScan.snp –-indel-file sample.VarScan.indel* –- *output-file sample.varScan.snp.filter*

(xi) java –jar VarScan.jar filter sample.VarScan.indel –-output-file sample.VarScan.indel.filter

(xii) java –jar VarScan.jar readcounts sample.mpileup.sam > sample.mpileup.readcounts

### Contamination check

Once the BAM files and IDs are generated, we could end up checking whether or not the BAM ids are error prone or contaminated across the samples using VerifyBAMID (Jun et al. 2012).

### (3) Downstream processing the files

VCF, an acronym for variant call format is a popular format to store variants calling data as it stores both SNPs and indel information succinctly. While BCF is a binary version of VCF, the format can be written and read using BCFtools tool using the following command:

(xiii) samtools mpileup -uf sample.sorted.bam | bcftools view - > sample.var.raw.bcf

While generating BCF file from BAM using samtools, -u is used for generating uncompressed VCF which can be piped as BCFtools designed for stream data and -f for the faidx-indexed reference file in the FASTA format.

(xiv) bcftools view sametools.var.raw.bcf | vcfutils.pl varFilter -D100 > sample.var.flt.vcf

(xv) samtools calmd -Abr sample.sorted.bam ~/hg38/hg38.fa > sample.baq.bam

(xvi) samtools mpileup -uf ~/hg38/hg38.fa sample.baq.bam | bcftools view - > sample.baq.var.raw.bcf

### Annotation and curation

Post processing the files, annotation and curation of the data followed by prioritizing the candidate SNPs/variants involves a great deal of user's discretion. There are a host of tools and annotation methods meant for this. Population stratification can be one such step in this process. While the 1000 genomes dataset(Auton et al. 2015) or gnomAD (Fu *et al. 2013*) containing the datasets are already used to summarize worldwide population, estimating individual ancestries using ADMIXTURE (Alexander et al. 2009) would help researchers to project the samples. On the other hand, downstream bioinformatics annotation can then be supplemented to integrate the results with different pathway tools, *viz*. PANTHER (Mi et al. 2016) which assesses the ontology/pathways affecting the “mutated” genes. This can be further supplemented by usage of assorted databases like Clinvar, dbSNP. In addition, global enrichment analysis and association networks using GeneMania (Warde-Farley et al. 2010) would allow create and visualize gene networks by evidence in pathways and gene-gene/protein-protein interactions (predicted and experimental).

### Benchmarking gave distinct distribution of variants

Benchmarking yielded a good recovery rate for validation of SNPs while VarScan with default values was found to have highest overall sensitivity with VarScan strict parameters having the lowest overall sensitivity (Figures 6 and 7). However, we observed that the preprocessing steps have little impact on the final output, with base recalibration step using GATK Unified Genotyper identifying fewer validated SNPs when compared to VarScan. On the other hand, we found that the recovery of exon variants among the exome samples was typically high when compared to the two whole genome datasets (Figure 5b). When variant lists were confined to previously observed variants as observed from the benchmark analyses between Sention and GATK (Weber et al. 2016), we observed that the recovery of SNPs with default parameter was found to be considerably good. Whereas changing variant calling criterion especially using VarScan, for example, imposing strict coverage requirement (Figure 7) yielded less numbers of false positives giving the number of *bona fide* or *de novo* variants (Figures 5a and 5b). This subtly proves that our benchmarking the six WES and two WGS datasets is variable with the capture, sequencing, processing and post-processing/analysis in the human genome and VarScan is comparable with the GATK in terms of identifying the *de novo* variants (Figure 5a and 5b). With the wet-lab components of NGS being cumbersome, analyzing the exons or for that matter intronic variants using bioinformatics pipeline is equally challenging. There must be significant *in silico* hurdles and organizational steps discussed from time to time and yet at the end of the analysis, one needs to arrive at the fittest in using the discretionary tools. Although technology challenges persist in setting up certain standards and guidelines, the end-user can enhance the pipeline with further tools. In this protocol, we have essentially shown how a WES pipeline can be run using batch file process and the comparison of VarScan over GATK using benchmarked datasets.

**Figure 3:**
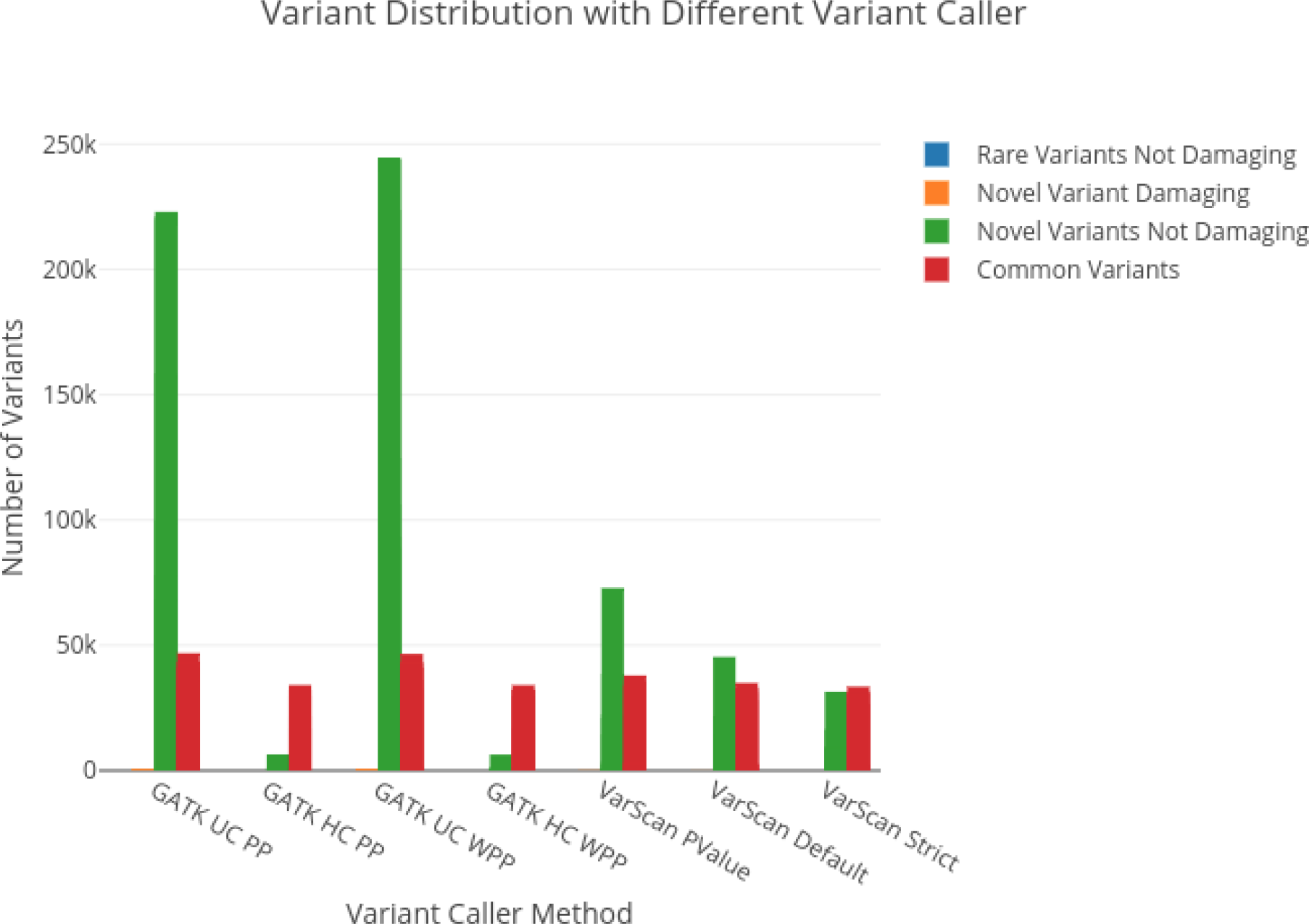
Number of variants obtained from GATK and VarScan with various parameters. We observe GATK Unified caller to have large number of false positives while VarScan with strict parameters performed well with less number of false positives.

**Figure 4:**
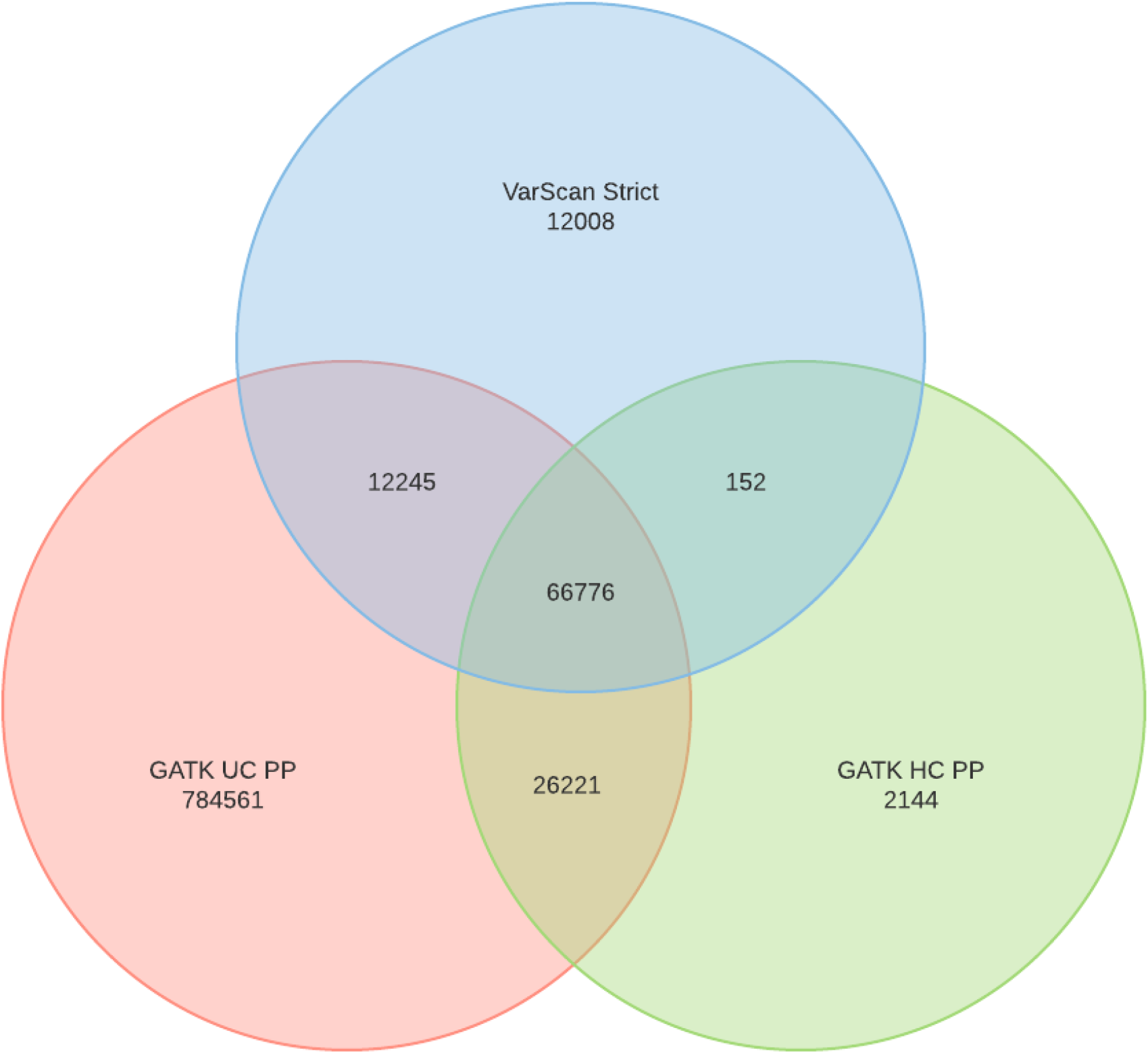
Venn diagram of three methods using Haplotype caller with preprocessing (HC-PP) and Universal genotype caller with preprocessing (UC-PP) and VarScan strict om sample SRR098359. We observed that all the three share the most true positive variants.

**Figure 5a:**
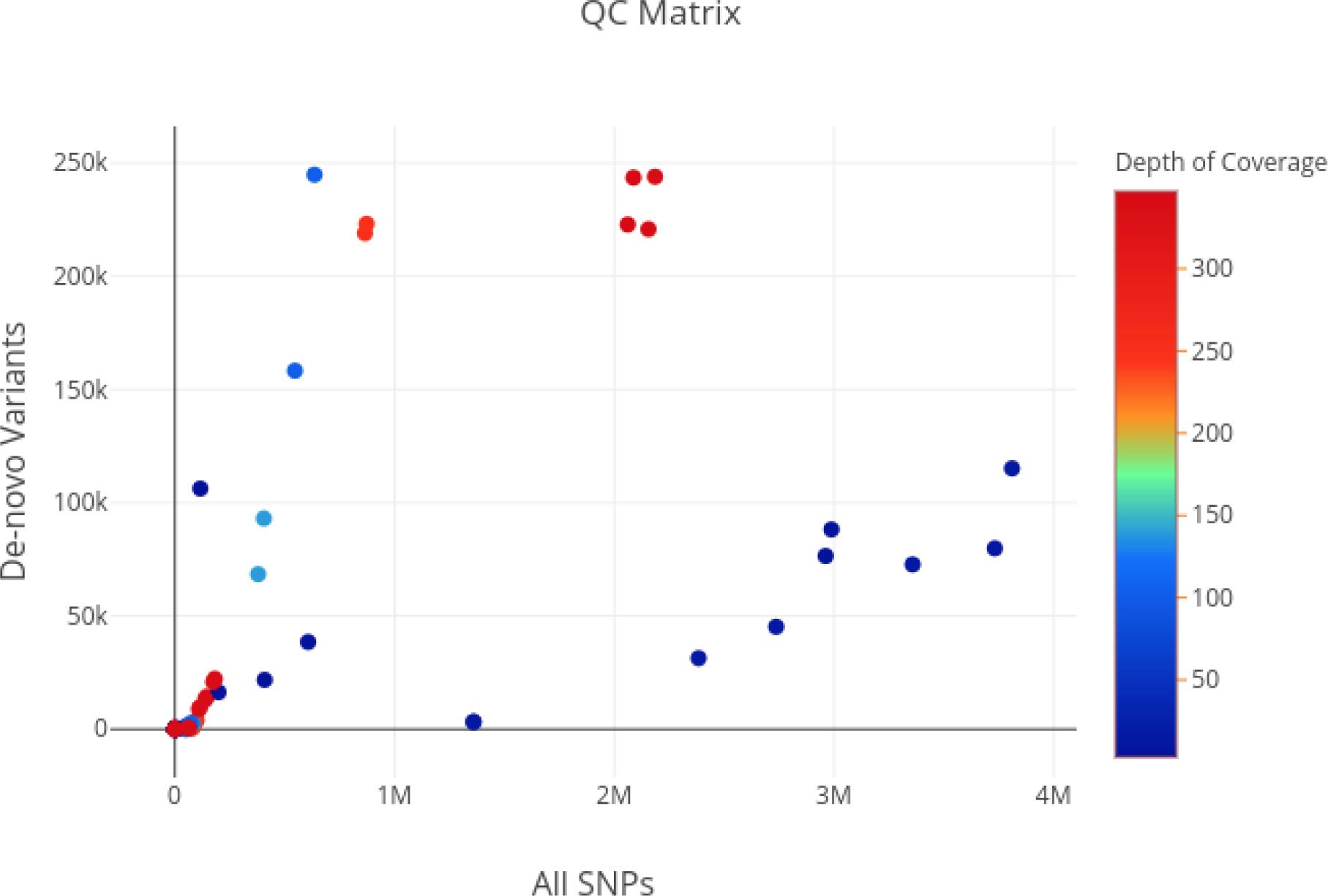
Distribution of *de novo* variants and all variants in regards to the depth of coverage of NGS run. The x-axis shows the number of million reads with depth of coverage shown inb legend (right) and the y-axis shows the number of *de novo* variants.

**Figure 5b:**
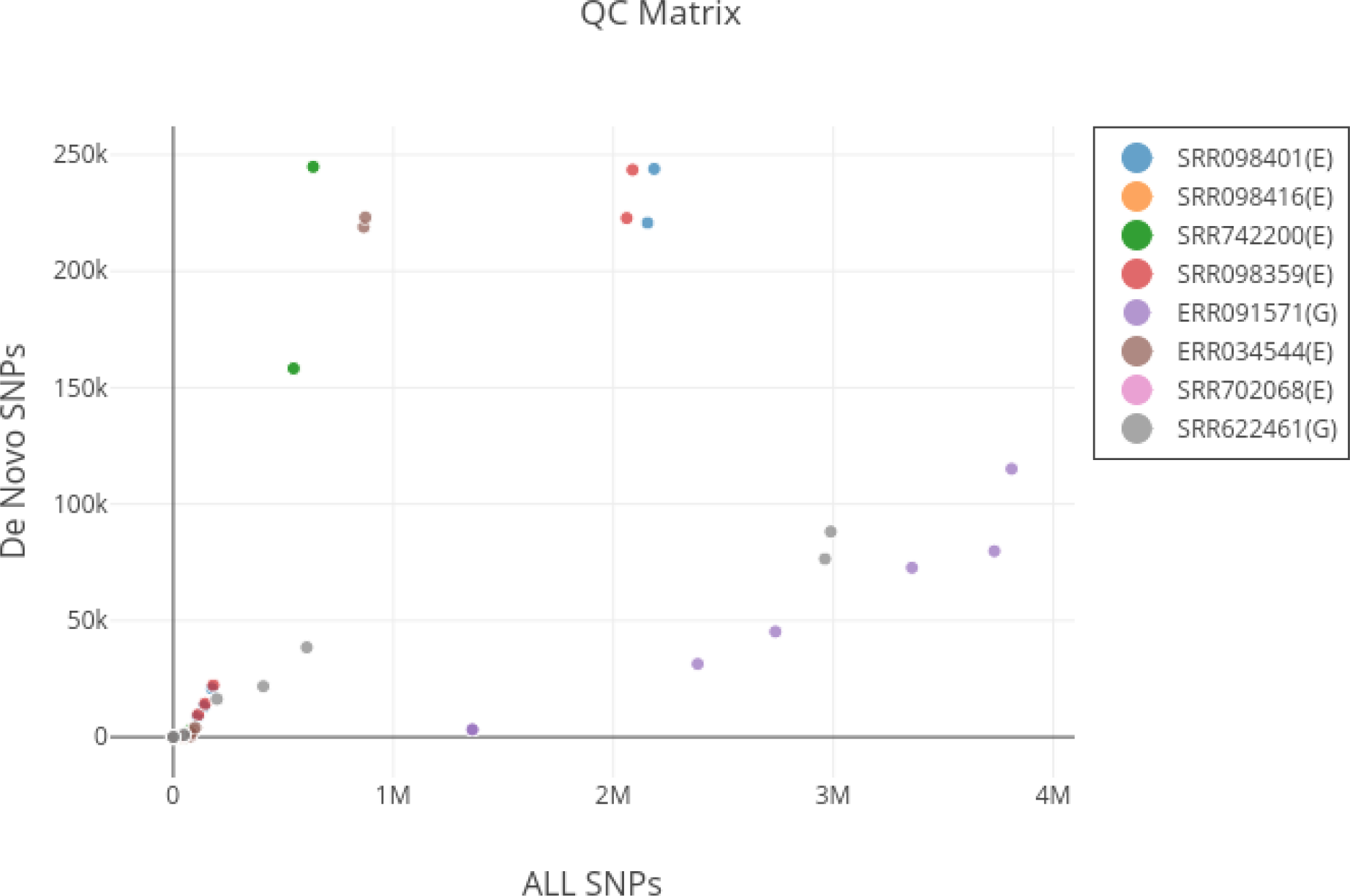
Distribution of *de novo* variants in regards to all SNPs against each sample.

**Figure 6:**
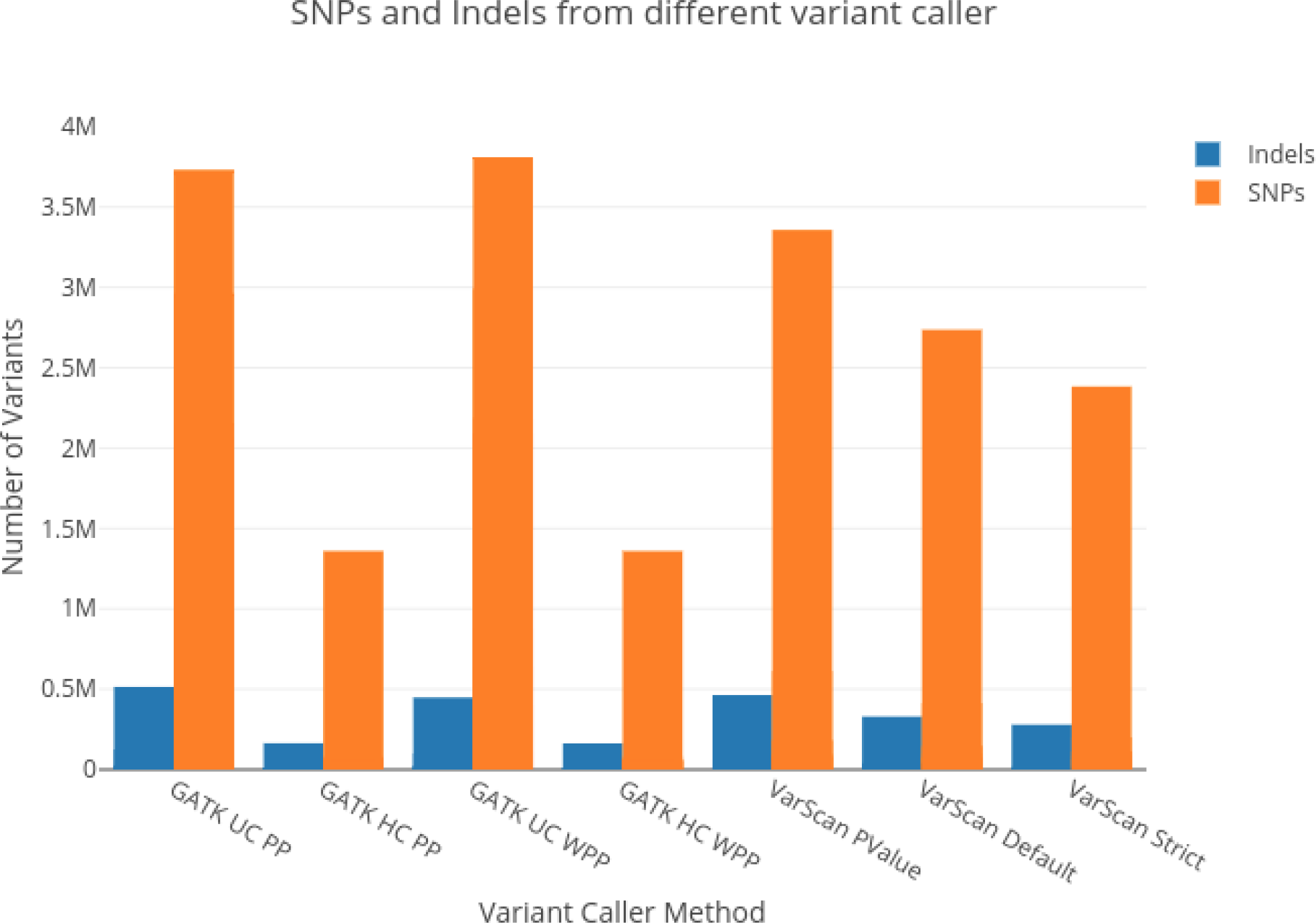
Number of SNPs and Indels called by GATK and VarScan using all parameters against the samples. We observed again that VarScan gave the best results with less false positive variants.

**Figure 7:**
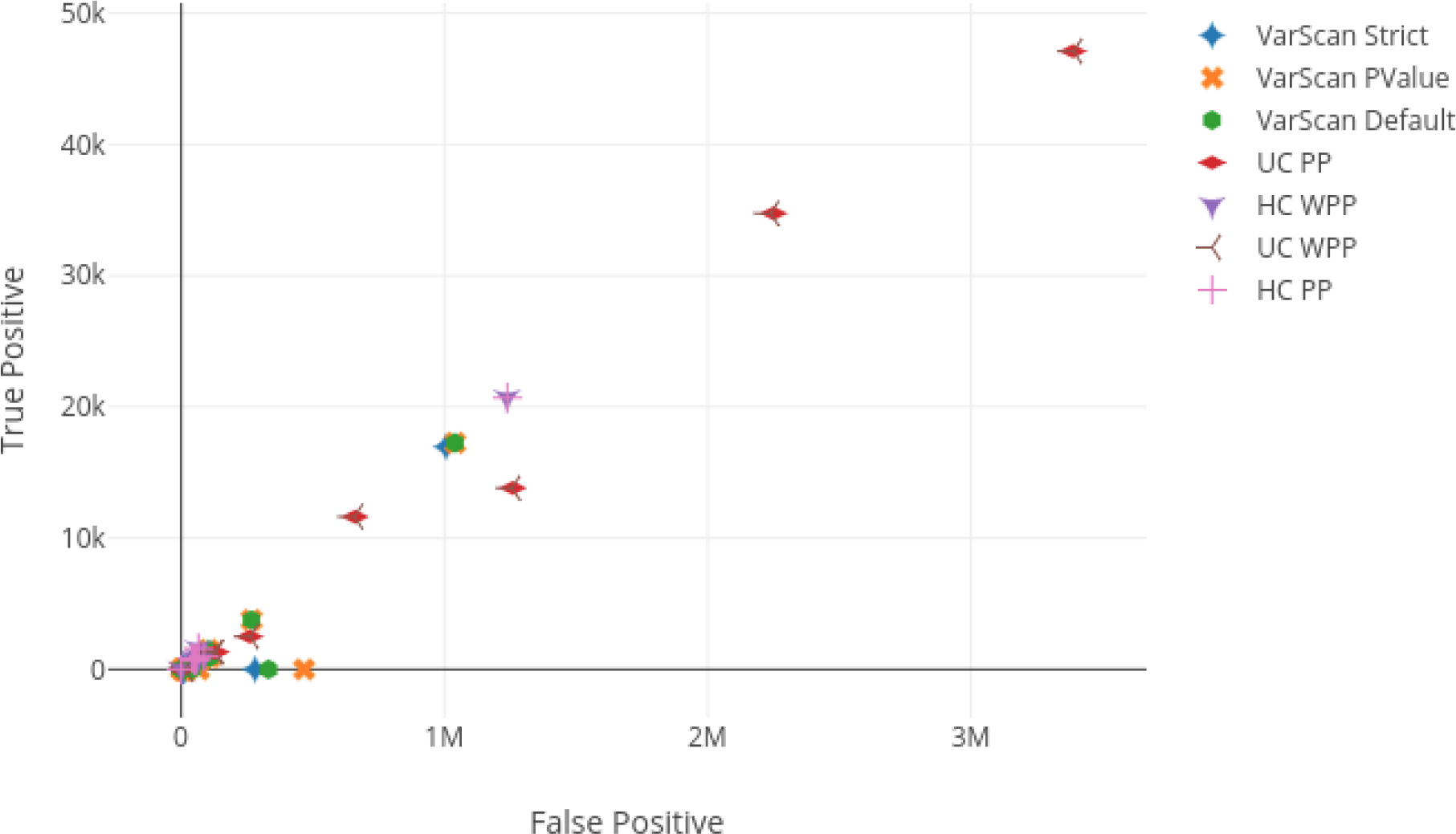
Scatter plot of number of true positives/false positives for all variant calling parameter options.

**Figure 8a:**
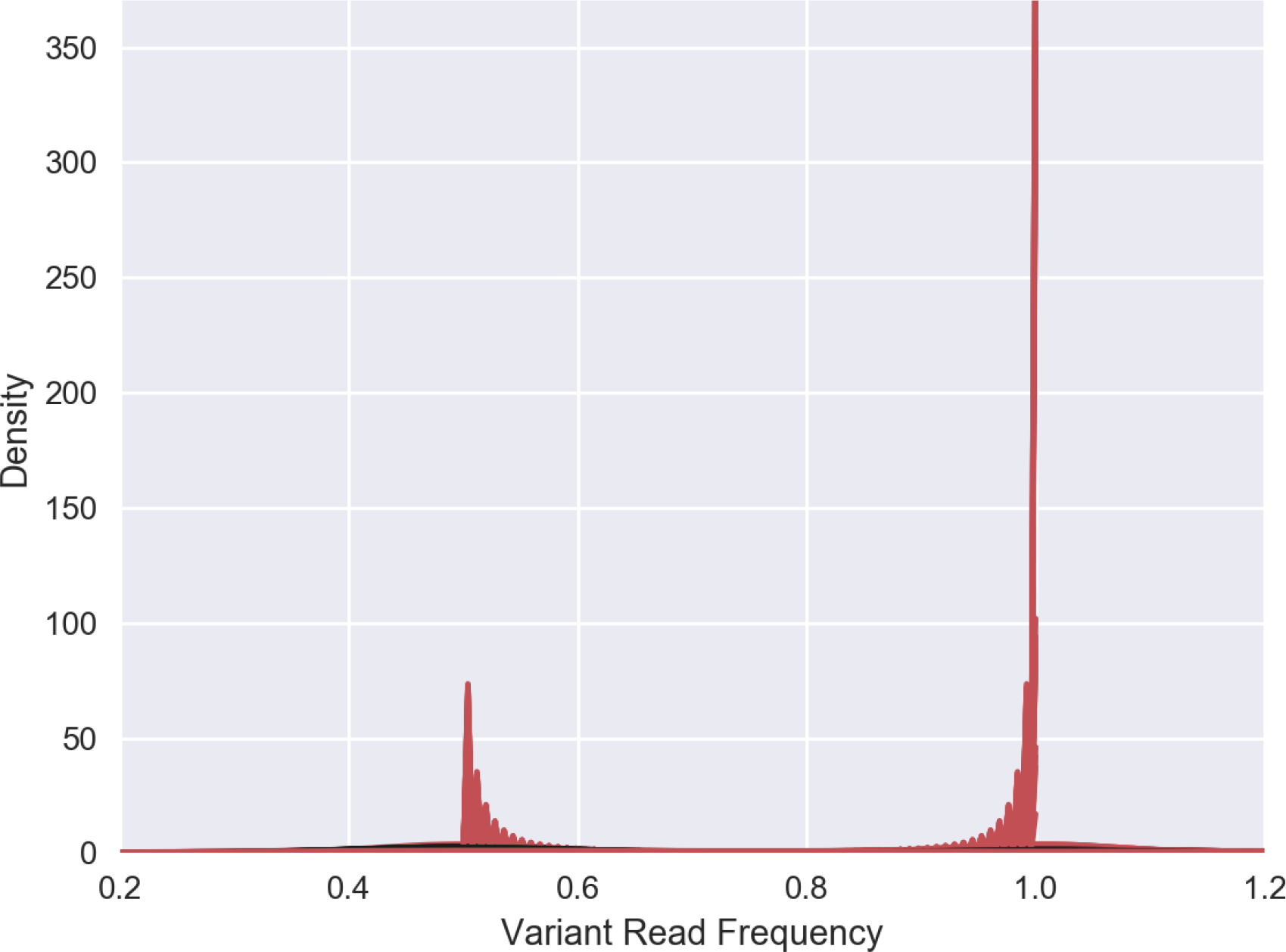
Density plot of an exome NGS run for *de novo* and known variants. The x-axis shows the variant read frequency against the density in y-axis.

**Figure 8b:**
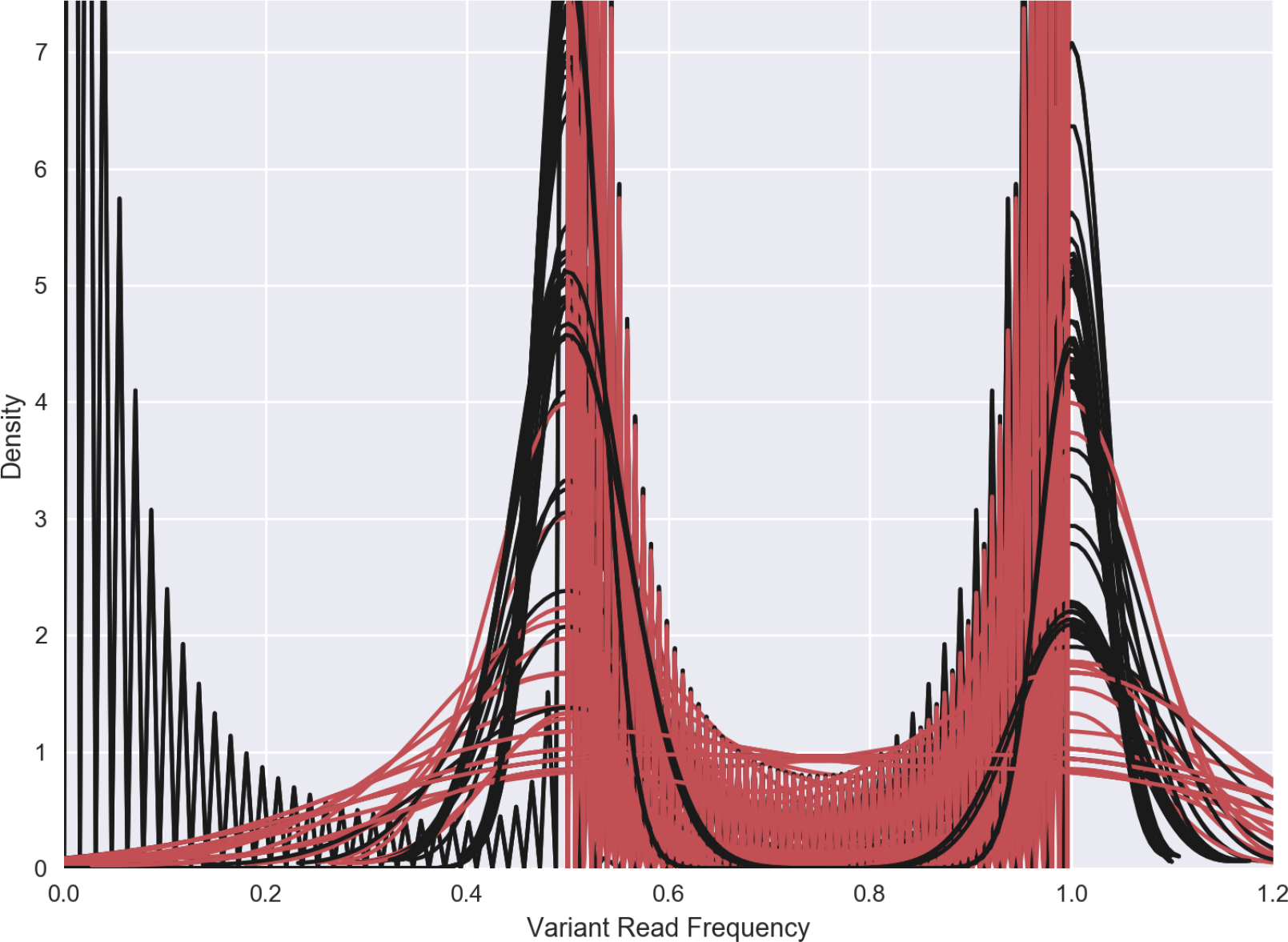
Density plot of NGS runs for *de novo* and known variants. The x-axis shows the variant read frequency against the density in y-axis.

**Table 2:** Variant calling pipelines and their respective arguments

| Pipeline | Arguments |
| --- | --- |
| VarScan-Default | Default |
| VarScan-Pvalue | --p-value 0.05 |
| VarScan Strict | --min-coverage 10 --min-avg-qual 20 or --min-reads2 4 --min-var-freq 0.3 |
| GATK WPP | "No Preprocessing" pipeline |
| GATK-PP | "Realign Only", "Recalibrate Only", and "Full Pipeline" |

## Acknowledgments

PS wants to acknowledge biostars.org forum which enabled him to enhance the pipeline consistently. He gratefully acknowledges the forum and immense discussions from users/researchers. The Birla Institute of Scientific Research would like to thank the Biotechnology Information System Network (BTIS), Department of Biotechnology, Government of India for funding and providing the resources and facilities. The authors gratefully acknowledge the Indian Council Medical research towards grant # 5/41/11/2012 RMC.

## Supplementary information

All the software can be downloaded/used from following locations:

1. FastQC https://www.bioinformatics.babraham.ac.uk/projects/fastqc/
2. Bowtie2 http://bowtie-bio.sourceforge.net/bowtie2/index.shtml
3. Samtools http://samtools.sourceforge.net/
4. VarScan http://varscan.sourceforge.net/
5. Bcftools https://github.com/samtools/bcftools
6. Vcftools https://github.com/samtools/bcftools
7. PANTHER http://pantherdb.org
8. dbSNP https://www.ncbi.nlm.nih.gov/projects/SNP/
9. 1000 genomes dataset http://www.internationalgenome.org/
10. GeneMania http://genemania.org/

